# Adaptive and Innate Immune Cell Responses in Tendons and Lymph Nodes After Tendon Injury and Repair

**DOI:** 10.1101/658310

**Authors:** Andrew C Noah, Thomas M Li, Leandro M Martinez, Susumu Wada, Jacob B Swanson, Nathaniel P Disser, Kristoffer B Sugg, Scott A Rodeo, Theresa T Lu, Christopher L Mendias

## Abstract

Tendon injuries are a common clinical condition with limited treatment options. The cellular components of the innate system, such as neutrophils and macrophages, have been well studied in tendon injuries. However the adaptive immune system, comprised of specialized lymphocytes, plays an important role in orchestrating the healing of numerous tissues but less is known about these cells in tendon healing. To gain a greater understanding of the biological processes that regulate tendon healing, we sought to determine how the cellular components of the adaptive and innate immune system respond to a tendon injury using two-month old male mice. We determined that the lymphatic vasculature is present in the epitenon and superficial regions of Achilles tendons. We then created an acute Achilles tenotomy followed by repair, and collected tendons and draining lymph nodes one, two, and four weeks after injury. Using flow cytometry and histology, after tendon injury we observed a robust adaptive immune cell response that followed an initial innate immune cell response. There was an accumulation of monocytes, neutrophils, and macrophages one week after injury that declined thereafter. Dendritic cells and CD4^+^ T cells peaked two weeks after injury, while B cells and CD8^+^ T cells progressively increased over time. In parallel, immune cells of the draining popliteal lymph node demonstrated a similarly coordinated response to the injury. These results suggest that there is an adaptive immune response to tendon injury and adaptive immune cells may play a role in regulating tendon healing.

## Introduction

Tendon injuries, which often occur due to excess mechanical loading, are a common clinical disorder in musculoskeletal medicine (20, 33). A scar is often formed at the site of tendon tears which can impair tissue biomechanical properties and overall mobility (13). The innate immune system, consisting in part of macrophages, neutrophils, and other related hematopoietic cell types, has been well studied in different models of tendon tear and repair, and is thought to play a role in the formation and remodeling of scar tissue (33). In acute tendon injuries, there is a rapid and transient accretion of neutrophils and pro-inflammatory M1 macrophages, followed by a slow and persistent accumulation of anti-inflammatory M2 macrophages (19, 30, 33). While numerous studies have documented the role of the innate immune system in tendon injuries, less is known about the potential roles of T cells, B cells, and other components of the adaptive immune system in tendon healing and scar formation.

Emerging evidence has indicated that both the adaptive and innate immune systems work along with tissue resident cells to coordinate tissue repair and fibrosis (12, 17). Upon peripheral tissue injury, signals such as cytokines and cells such as dendritic cells bearing antigens from the injured site travel via lymphatic vessels to the draining lymph node to stimulate the T and B cells there to initiate an adaptive immune response (3, 17). With immune responses, the draining lymph node undergoes hypertrophy, reflecting both increased lymphocyte trafficking to and proliferation in lymph nodes (3). While recent studies in humans have identified that the adaptive immune system is activated in chronic degenerative tendinopathies (21), less is known about whether the adaptive immune response plays a role in frank tendon tears. Additionally, the lymphatic system which allows for the recovery of interstitial fluid from tissue and regulates the immune cell response to an injury (3, 27), has not been well studied in the context of tendon injuries. Since further understanding of the role of the adaptive immune response may help to identify potential therapeutic interventions to improve outcomes for patients with tendon injuries, our objective was to study the adaptive immune cell and lymph node response to an acute tendon injury. We hypothesized that an adaptive immune cell response would be present after an acute tendon injury, along with a corresponding lymph node response. To test this hypothesis, we identified the location of the lymphatic vasculature in Achilles tendons and draining lymph nodes, and then performed an acute Achilles tenotomy and repair in mice. Tendons and lymph nodes were collected one, two, and four weeks after the surgical intervention. Flow cytometry and immunofluorescence staining of tissue sections were performed using markers of innate and adaptive immune cells in parallel with measurement of changes in expression of genes with known roles in immune cell recruitment, extracellular matrix (ECM) composition, and tenogenesis.

## Methods

### Animals

This study was approved by the Hospital for Special Surgery/Weill Cornell Medical College/Memorial Sloan Kettering Cancer Center IACUC (protocol 2017-0035). All experiments were conducted in accordance with our IACUC protocol and were consistent with the US Public Health Service Policy on the Humane Care and Use of Laboratory Animals. Two-month old male C57BL/6J mice (strain 000664, Jackson Labs, Bar Harbor, ME) and transgenic mice that express GFP under the control of a 4kb segment of the scleraxis promoter (ScxGFP (24)) were used in this study. Mice were housed under specific pathogen free conditions and provided with *ad libidum* access to food and water.

### Surgeries

To identify the lymph node that drains the Achilles tendon, mice (N=4) were deeply anesthetized with 2% isoflurane, and the skin overlying the lower limb was shaved and cleaned with 4% chlorhexidine. Approximately 20μL of 2% Evan’s Blue Dye (EBD, Sigma Aldrich, St Louis, MO) was injected into the peritendinous space of Achilles tendons of mice. The EBD was allowed to drain into the lymphatics and draining lymph node for fifteen minutes, and mice were then euthanized with carbon dioxide followed by cervical dislocation. The skin overlying the lower limb was carefully dissected from the mice to visualize the popliteal lymph nodes.

A separate cohort of mice underwent an Achilles tenotomy and repair, using techniques modified from a previous study (23, 30). Briefly, mice were deeply anesthetized with 2% isoflurane and placed in a prone position. The skin overlying the surgical site was shaved and cleaned with 4% chlorhexidine. A midline skin incision was made on the posterodistal aspect of the hindlimb and the paratenon was reflected. A full-thickness tenotomy was completed in the mid-substance of the Achilles tendon followed by immediate two-strand repair using a Modified Kessler technique with 6-0 Prolene suture (Ethicon, Somerville, NJ). The skin was then closed with interrupted sutures using 6-0 Prolene suture (Ethicon). The plantaris tendon was left intact. Buprenorphine (0.05mg/kg, Reckitt, Parsippany, NJ) was administered for analgesia during the postoperative recovery period. Weightbearing and cage activity were allowed after surgery, and mice were closely monitored thereafter for signs of pain or distress. Surgical interventions were well tolerated.

At either one, two, or four weeks after tenotomy and repair (N=12 mice per time point), mice were sacrificed by exposure to carbon dioxide followed by cervical dislocation, and Achilles tendons were harvested for flow cytometry, gene expression, and histology. Popliteal lymph nodes were also harvested for flow cytometry. Achilles tendons and popliteal lymph nodes were also collected from control mice that did not undergo surgical repair. The Achilles tendons of an additional set of uninjured ScxGFP mice (N=4) were collected as described above for experiments to visualize the presence of the lymphatic vasculature in tendons. Bilateral tissues were combined for flow cytometry and gene expression analysis.

### Histology

Histology was performed as previously described (9, 15). Achilles tendons were placed in 30% sucrose solution for one hour, and then snap frozen in Tissue-Tek OCT compound (Sakura Finetek, Torrance, CA) and stored at −80°C until use. Tissues were sectioned at a thickness of 10μm in a cryostat. For the immunolocalization of the lymphatic vasculature in tendon, slides were fixed in 4% paraformaldehyde, permeabilized in 0.2% Triton X-100 and blocked with 5% goat serum. Slides were incubated with primary antibodies against CD31 (1:100, catalog number 550274, BD, San Diego, CA) to label endothelial cells, and LYVE-1 (1:200, catalog number 14044380, Thermo Fisher Scientific) and podoplanin (1:300, catalog number 127408, BioLegend, San Diego, CA) to label the lymphatic vasculature (14). To visualize the immune cell response after injury in tendon, tissues were fixed in ice cold acetone for ten minutes, blocked with 5% goat serum for one hour, and stained with biotinylated antibodies against CD11b to identify myeloid cells (1:1000, catalog number 101204, BioLegend), CD3 to identify T cells (1:250, catalog number ab5690, Abcam) and FITC conjugated antibodies against B220 to identify B cells (1:100, catalog number 103206, BioLegend). Secondary antibodies conjugated to AlexaFluor 405 (1:400, catalog number ab175671, Abcam), AlexaFluor 555 (1:400, catalog number A21428, Themo Fisher Scientific), or AlexaFluor 647 (1:400, catalog number 405510, BioLegend; 1:500, catalog number A21245, Thermo Fisher Scientific) or Streptavidin conjugated to AlexaFluor 647 (1:500, S32357, Invitrogen) were used to detect primary antibodies that were not already conjugated to a fluorophore. Nuclei were stained with DAPI (1:500, Sigma Aldrich, St. Louis, MO). High-resolution images were captured with an Eclipse Ni-E microscope (Nikon, Melville, NY).

### Flow Cytometry

Freshly harvested tendons and lymph nodes were prepared for flow cytometry as modified from previous studies (15, 28). Tissues were finely minced, and then digested in type II collagenase (616 U/mL, Worthington Biochemical, Lakewood, NJ), DNase I (80μg/mL, Sigma Aldrich), and EDTA (20μL/mL, Thermo Fisher Scientific) for 40 minutes at 37°C. Cell suspensions were subsequently triturated with glass pipettes and passed through 70μm filters to remove undigested debris. Total cell counts were determined using a Z Series Coulter Counter (Beckman Coulter, Indianapolis, IN).

Cells were labeled with fluorophore-conjugated antibodies from BioLegend, including antibodies against B220 (catalog number 103206), CD3 (catalog number 100330), CD4 (catalog number 100434), CD8 (catalog number 100730), CD11b (catalog number 101233), CD11c (catalog number 117324), CD64 (catalog number 139304), Ly6C (catalog number 128014), and MHC II (catalog number 116415). Cells were then fixed and permeabilized with Cytofix (BD) and then labeled with antibodies against intracellular IgG (catalog number 405308) to detect intracellular IgG that are high in plasma cells (15). Gating strategies are based on our previously published work (1, 11, 15). While the use of intracellular staining precluded the use of DAPI to assess dead cells, parallel samples that were not fixed, but were labeled with CD45 (catalog number 103127) and DAPI (catalog number 422801) demonstrated that 95% of tendon and lymph node CD45+ cells were DAPI^−^, indicating that they were viable cells. Cells were analyzed using a FACSCanto (BD) and FlowJo software (version 10, Tree Star, Ashland, OR).

### Gene Expression

RNA isolation and gene expression was performed as modified from previous reports (5, 7, 31). Achilles tendons were placed in TRIzol Reagent (Life Technologies, Carlsbad, CA) in tubes containing 1.5mm zirconium beads (Benchmark Scientific, Sayreville, NJ), processed in a shaking bead mill for two cycles of 3 minutes, and then snap frozen and stored at −80°C. Samples were thawed on ice, and RNA was isolated using MaXtract High Density columns (Qiagen, Valencia, CA), followed by further purification with Zymogen IC spin columns (Zymo Research, Irvine, CA). RNA was treated with DNase I (Qiagen), and reverse transcribed into cDNA with iScript Reverse Transcription Supermix (Bio-Rad, Hercules, CA). Amplification of cDNA was performed in a CFX96 real-time thermal cycler (Bio-Rad) using SsoFast Universal SYBR Green Supermix (Bio-Rad). Target gene expression was normalized to the stable housekeeping gene *Ppid* using the 2^-ΔCt^ method. Primer sequences are provided in Table 1.

**Table 1.**
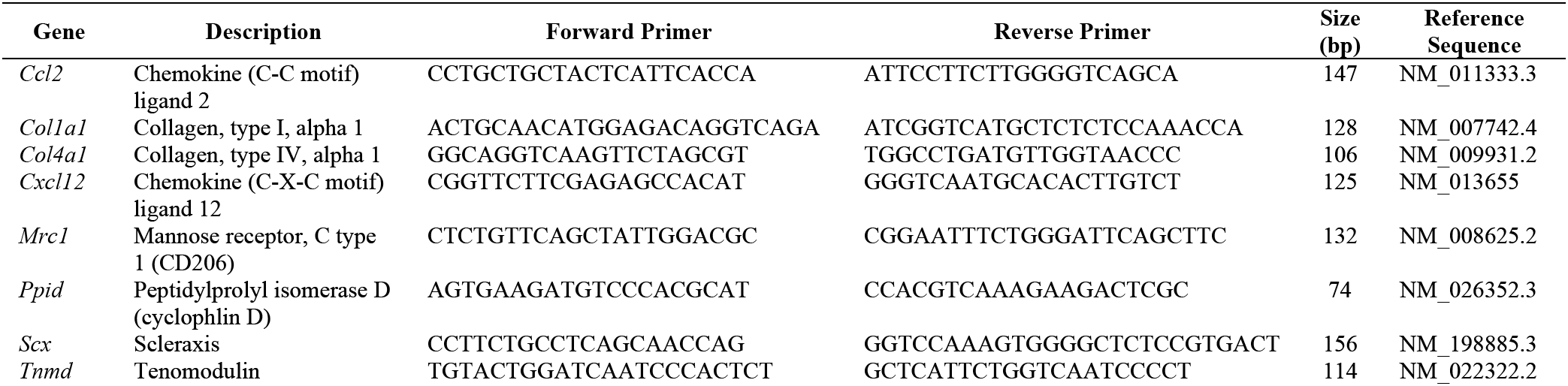
Primer sequences used for quantitative PCR.

### Statistics

Data is presented as mean±SD. For normally distributed data, differences between groups were tested using a one-way ANOVA followed by Bonferroni post-hoc sorting. Data that was not normally distributed was tested using a Kruskal-Wallis test followed by Dunn’s multiple comparisons test. Statistical analyses were conducted using Prism software (version 8.0, GraphPad, La Jolla, CA). Differences were considered statistically significant when P<0.05.

## Results

We first sought to confirm that the Achilles tendon and surrounding tissue drained to the popliteal lymph node. EBD was injected into the peritendinous space of the Achilles tendon, and fifteen minutes later a visible collection of EBD was found in the popliteal lymph node (Figure 1A). After determining that the Achilles tendon drains to the popliteal lymph node, we identified the location of the lymphatic vasculature in Achilles tendons using ScxGFP mice, which express GFP in scleraxis-expressing tenocytes. We observed lymphatic vessels, which jointly express the endothelial cell marker CD31, the lymphatic endothelial cell marker podoplanin, and lymphatic capillary marker LYVE1 in the epitenon that surrounds Achilles tendons (Figures 1B-C). The lymphatic vessels that enter into the superficial tendon were flanked by scleraxis-expressing tenocytes (Figure 1C). The lymphatic vessels were also in close proximity to the blood vasculature, as identified by CD31 vessels that did not contain LYVE1 or podoplanin (Figure 1C).

**Figure 1.**
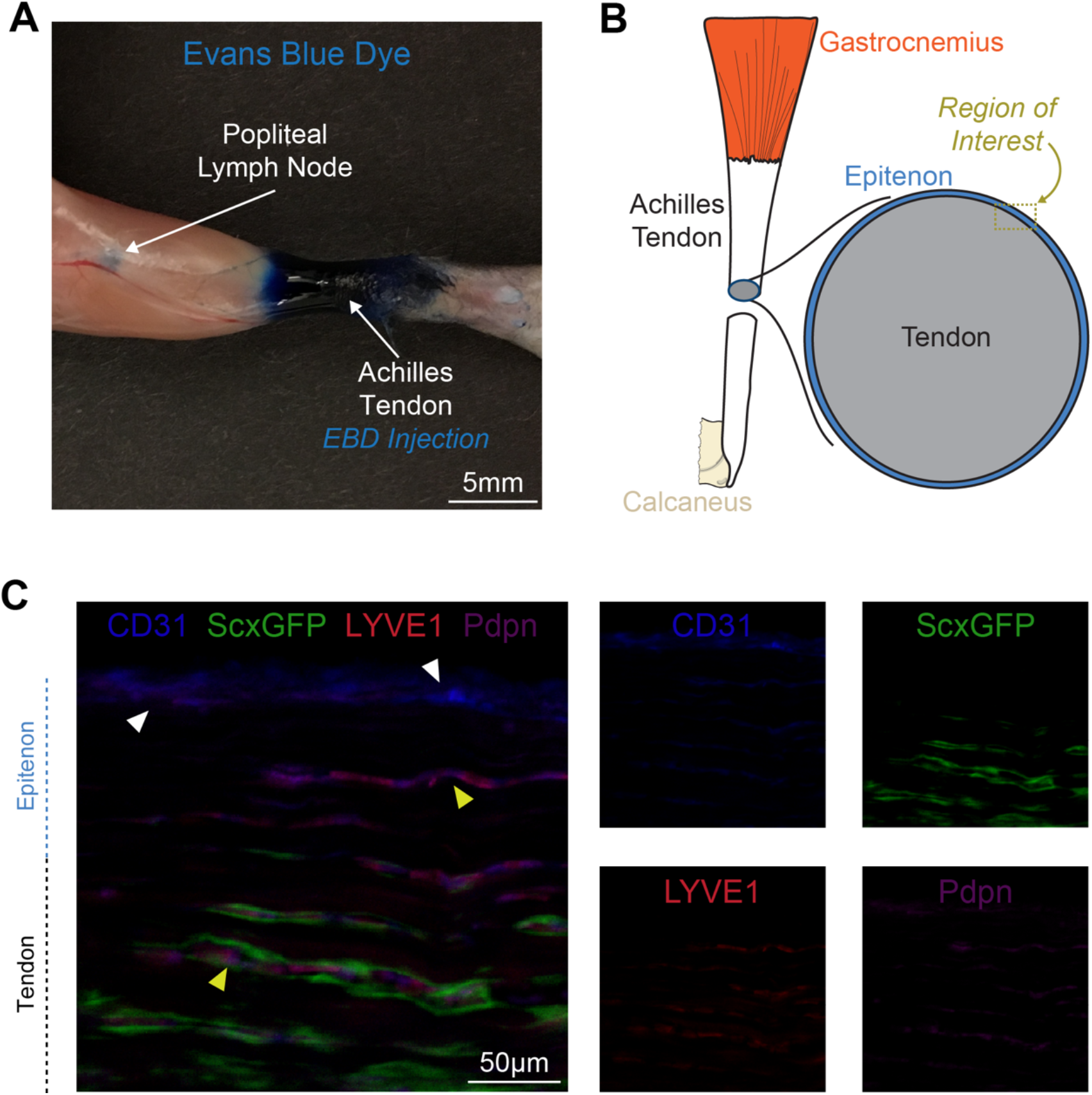
Lymph node drainage and lymphatic vessel location in tendon. (A) Posterior view of the lower limb demonstrating that 15 minutes after Evans Blue Dye (EBD) injection into the Achilles peritendinous region there is an accumulation of EBD in the popliteal lymph node. (B) Overview of the region of interest for panel C, which contains the epitenon and superficial tendon regions. (C) Histology demonstrating the presence of blood vasculature as indicated by CD31 signal in the absence of podoplanin and LYVE1 (white arrowheads), and lymphatic vasculature (yellow arrowheads) as indicated by CD31 with podoplanin and LYVE1 signal, with adjacent scleraxis-expressing tenocytes. Each channel is also shown individually in smaller panels. Blue, CD31; red, LYVE1; magenta, podoplanin; green, scleraxis-GFP. Scale bar is 50μm.

Following the studies that identified the lymphatic drainage from the Achilles tendon, we then determined innate and adaptive immune cell populations in tendons through four weeks after tenotomy and repair. There was an initial upregulation in the ECM genes *Col1a1* and *Col4a1* one week after tenotomy and repair, with a peak observed at two weeks (Figure 2). The tenogenic marker genes *Scx* and *Tnmd* were elevated beginning at two weeks after injury (Figure 2). There was also an initial upregulation in the monocyte chemoattractant gene *Ccl2* at one week, while *Cxcl12,* which can promote the trafficking of multiple immune cell populations including T-and B cells, was not upregulated until two weeks after injury (Figure 2). The M2 macrophage marker, CD206, peaked at two weeks and remained elevated at four weeks (Figure 2).

**Figure 2.**
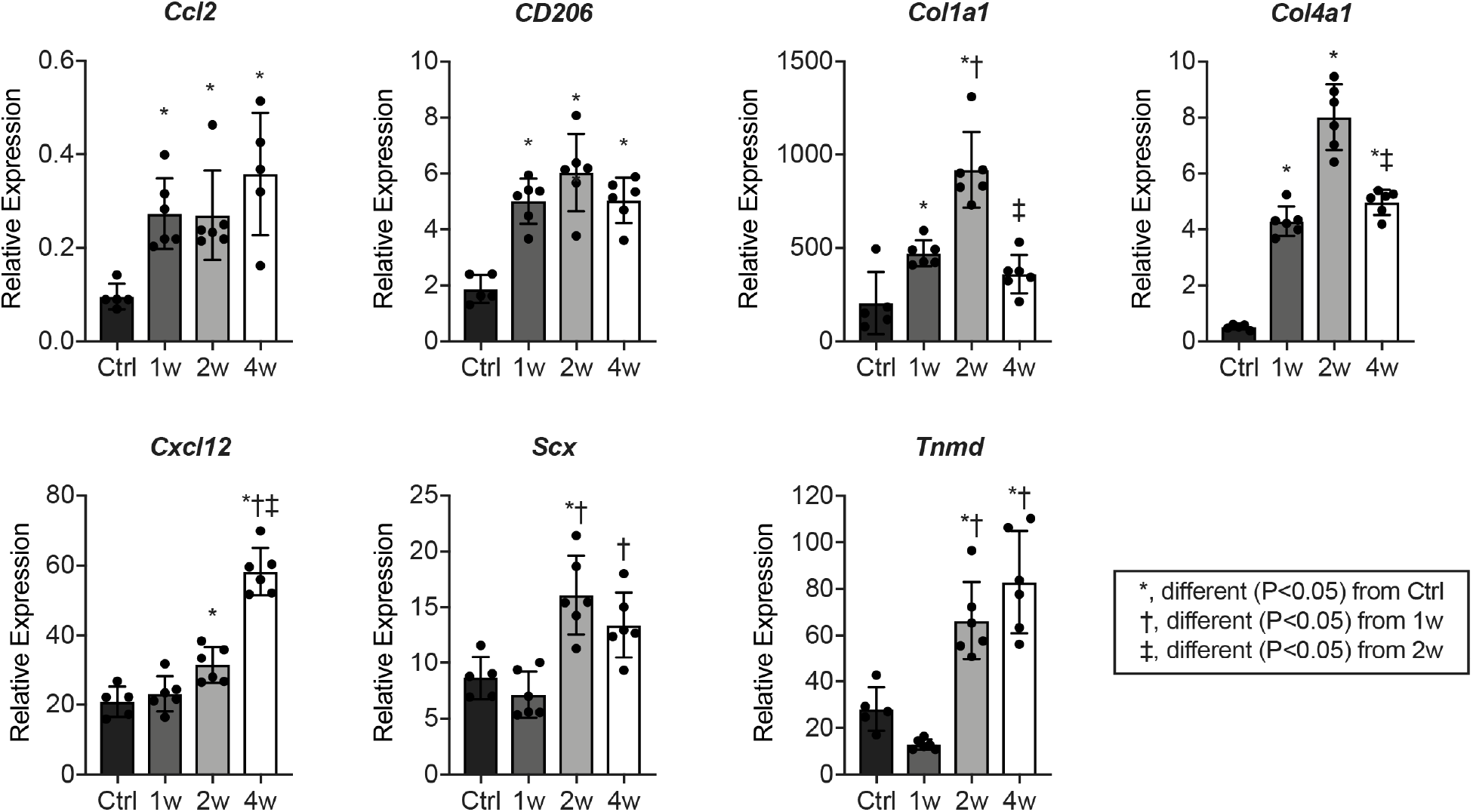
Expression of genes involved in immune cell recruitment, extracellular matrix composition, and tenogenesis after Achilles tenotomy and repair. Values are presented as mean±SD. Differences between groups tested using a one-way ANOVA (α=0.05) followed by Bonferroni post-hoc sorting: *, significantly different (P<0.05) from control (Ctrl); †, significantly different (P<0.05) from 1 week (1w); ‡, significantly different (P<0.05) from 2 week (2w). N≥3 mice per time point.

Consistent with the idea of early recruitment of innate cells, there was a marked accumulation of CD11b^+^/Ly6C^hi^ monocytes and CD11b^+^/Ly6C^med^/SSC^hi^ neutrophils at one week after injury, followed by their precipitous decline (Figure 3A-B). CD11b^+^/Ly6C^lo^/CD64^+^ macrophages were similarly transiently increased (Figure 3A-B), presumably reflecting both the differentiation of accumulated monocyte-derived macrophages and the proliferation of resident macrophages. The initial accumulation of these innate cells was followed by an accumulation of CD4^+^ T cells (CD3^+^/CD4^+^/CD8^−^) and CD11c^+^ MHC II^+^ antigen-presenting cells likely composed mostly of dendritic cells at two weeks after injury, while B cells (CD11b^−^/CD11c^−^ /B220^+^) and CD8^+^ T cells (CD3^+^/CD4^−^/YCD8^+^) demonstrated a progressive increase over time (Figures 3A-B). We observed similar qualitative results in histology experiments (Figure 4A) evaluating the presence of innate (Figure 4B) and adaptive (Figure 4C) markers in injured tendons.

**Figure 3.**
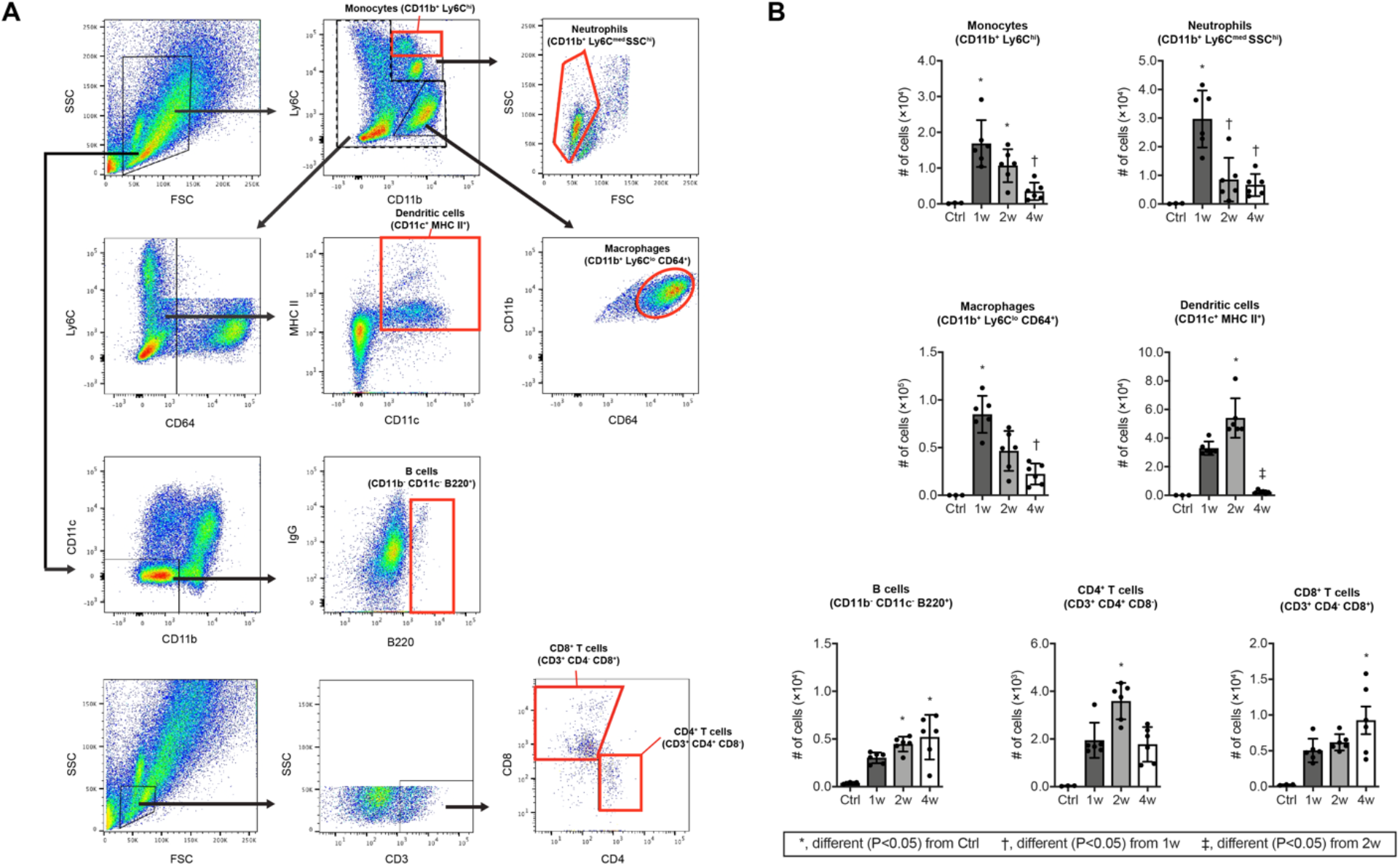
Flow cytometry of immune cell populations in Achilles tendons after tenotomy and repair. (A) Dot plot demonstrating gating strategies and (B) quantification of populations of monocytes (CD11b^+^/Ly6C^hi^), neutrophils (CD11b^+^/Ly6C^med^/SSC^hi^), macrophages (CD11b^+^/Ly6C^lo^/CD64^+^), dendritic cells (CD11c^+^/MHC II^+^), B cells (CD11bYCD11cYB220^+^), CD4^+^ T cells (CD3^+^/CD4^+^/CD8^−^), and CD8^+^ T cells (CD3^+^/CD4^−^/CD8^+^) from Achilles tendons. Differences between groups tested using a one-way ANOVA (α=0.05) followed by Bonferroni post-hoc sorting, as well as Kruskal-Wallis test followed by Dunn’s multiple comparisons: *, significantly different (P<0.05) from control (Ctrl); †, significantly different (P<0.05) from 1 week (1w); ‡, significantly different (P<0.05) from 2 week (2w). N≥3 mice per time point.

**Figure 4.**
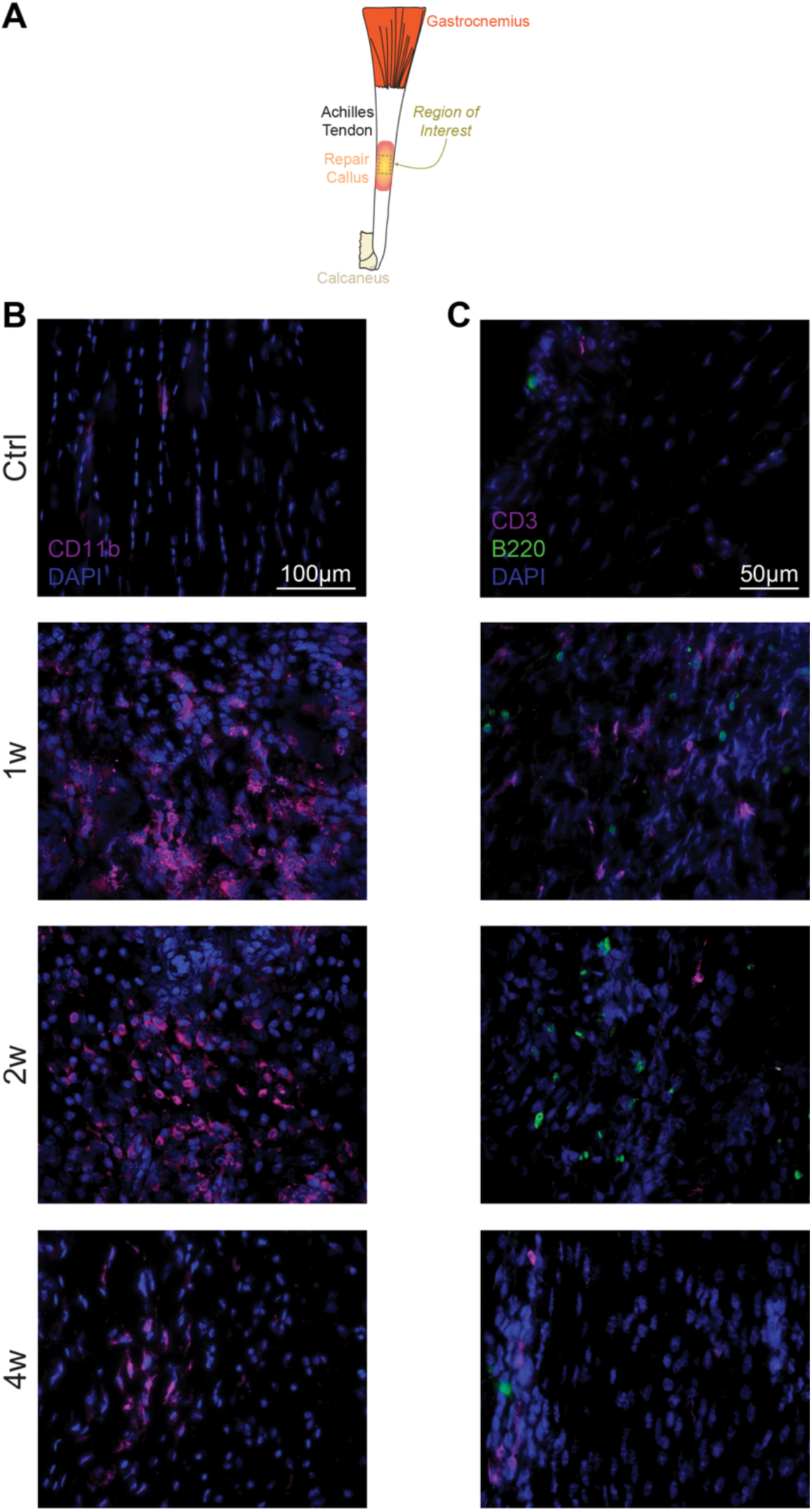
Location of immune cells in injured tendons. (A) Overview of the region of interest for panels B and C, which is taken from the midsubtance of the Achilles tendon. In the groups that underwent a tenotomy and repair, this region was in the callus that formed in the repaired area. (B) Myeloid cells, as indicated by CD11b signal, in control tendons, and tendons either 1 week (1w), 2 weeks (2w), or 4 weeks (4w) after tenotomy and repair. Blue, nuclei (DAPI); magenta, CD11b. Scale bar for all panels is 100μm. (C) T cells and B cells, as indicated by CD3 and B220 signal, respectively, in control tendons, and tendons either 1w, 2w, or 4w after tenotomy and repair. Blue, nuclei (DAPI); magenta, CD3; green, B220. Scale bar for all panels is 50μm.

Finally, we evaluated changes in innate and adaptive cell populations of popliteal lymph nodes after tenotomy and repair, and observed lymph node hypertrophy with a similar pattern of innate cell accumulation followed by adaptive cell accumulation. There was a pronounced accumulation of monocytes and macrophages one week after injury, while no differences in neutrophils or plasma cells (CD11b^−^/CD11c^−^/B220^−^/IgG^+^) were observed (Figures 5A-B). Migratory (CD11c^med^/MHC II^hi^) and resident (CD11c^hi^/MHC II^med^) dendritic cells showed a more gradual accumulation, starting at one week and peaking at two weeks. The bulk of the lymph node hypertrophy reflected the accumulation of CD4^+^ and CD8^+^ T cells and B cells, which reached plateau numbers by one week but were maintained through the second week (Figures 5A-B). Plasma cells, the differentiated effector B cells that secrete antibodies, were not statistically different across time points but showed a trend to increase at four weeks (p=0.058).

**Figure 5.**
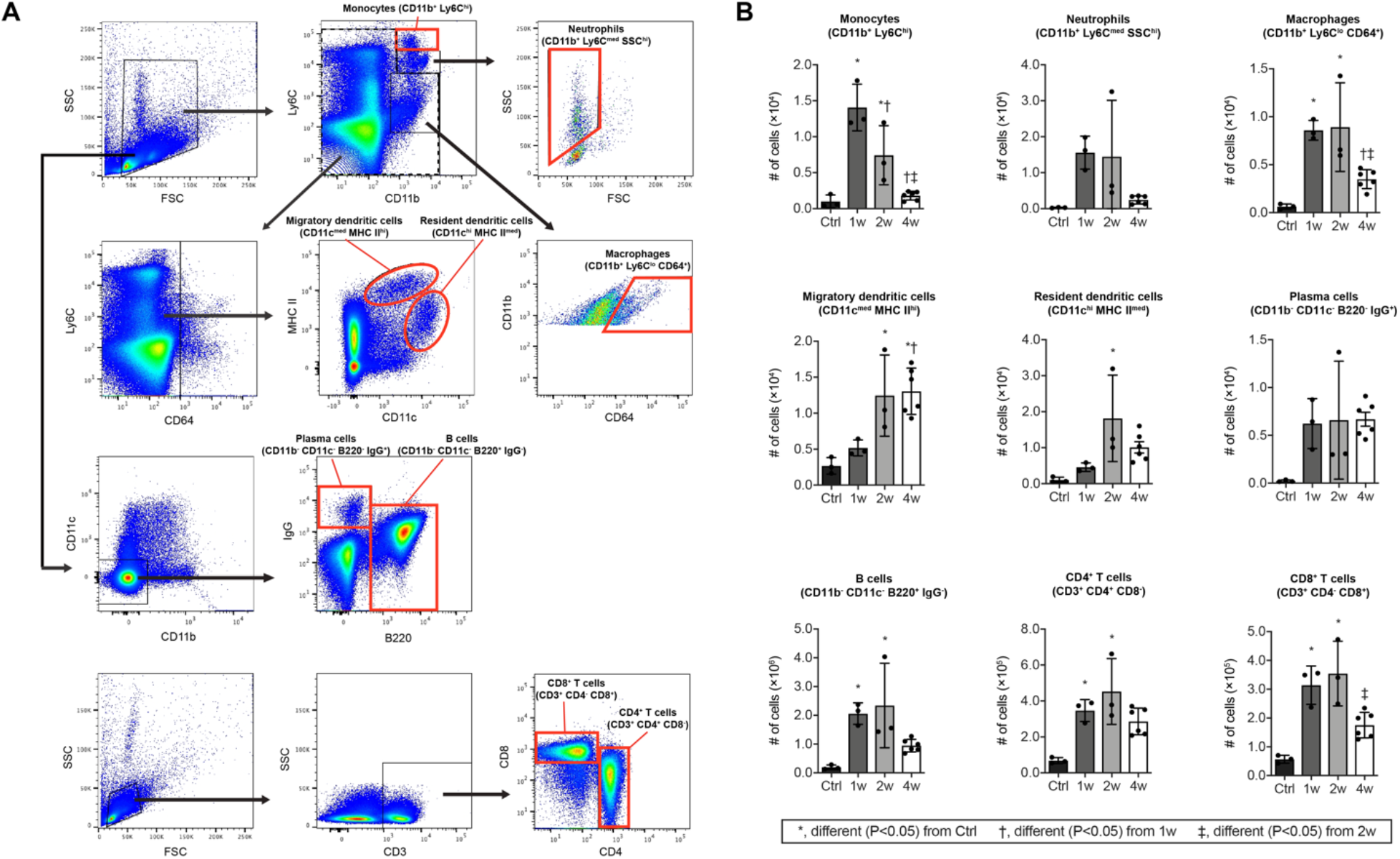
Flow cytometry of immune cell populations in popliteal lymph nodes after Achilles tenotomy and repair. (A) Dot plot demonstrating gating strategies and (B) quantification of populations of monocytes (CD11b^+^/Ly6C^hi^), neutrophils (CD11b^+^/Ly6C^med^/SSC^hi^), macrophages (CD11b^+^/Ly6C^lo^/CD64^+^), migratory dendritic cells (CD11c^med^/MHC II^hi^), resident dendritic cells (CD11c^hi^/MHC II^med^), plasma cells (CD11b^−^/CD11c^−^/B220/^−^IgG^+^), B cells (CD^−^/CD11c^−^ /B220^+^/IgG^−^), CD4^+^ T cells (CD3^+^/CD4^−^/CD8^−^), and CD8^+^ T cells (CD3^+^/CD4YCD8^+^) from popliteal lymph nodes. Differences between groups tested using a one-way ANOVA (α=0.05) followed by Bonferroni post-hoc sorting, as well as Kruskal-Wallis test followed by Dunn’s multiple comparisons: *, significantly different (P<0.05) from control (Ctrl); †, significantly different (P<0.05) from 1 week (1w); ‡, significantly different (P<0.05) from 2 week (2w). N≥3 mice per time point.

The lymph node hypertrophy and presence of plasma cells support the idea that the draining lymph node is responding to the tendon injury.

In addition to the individual cell populations presented in Figures 3B and 5B, Figure 6 provides a summary of major cell types across all time points.

**Figure 6.**
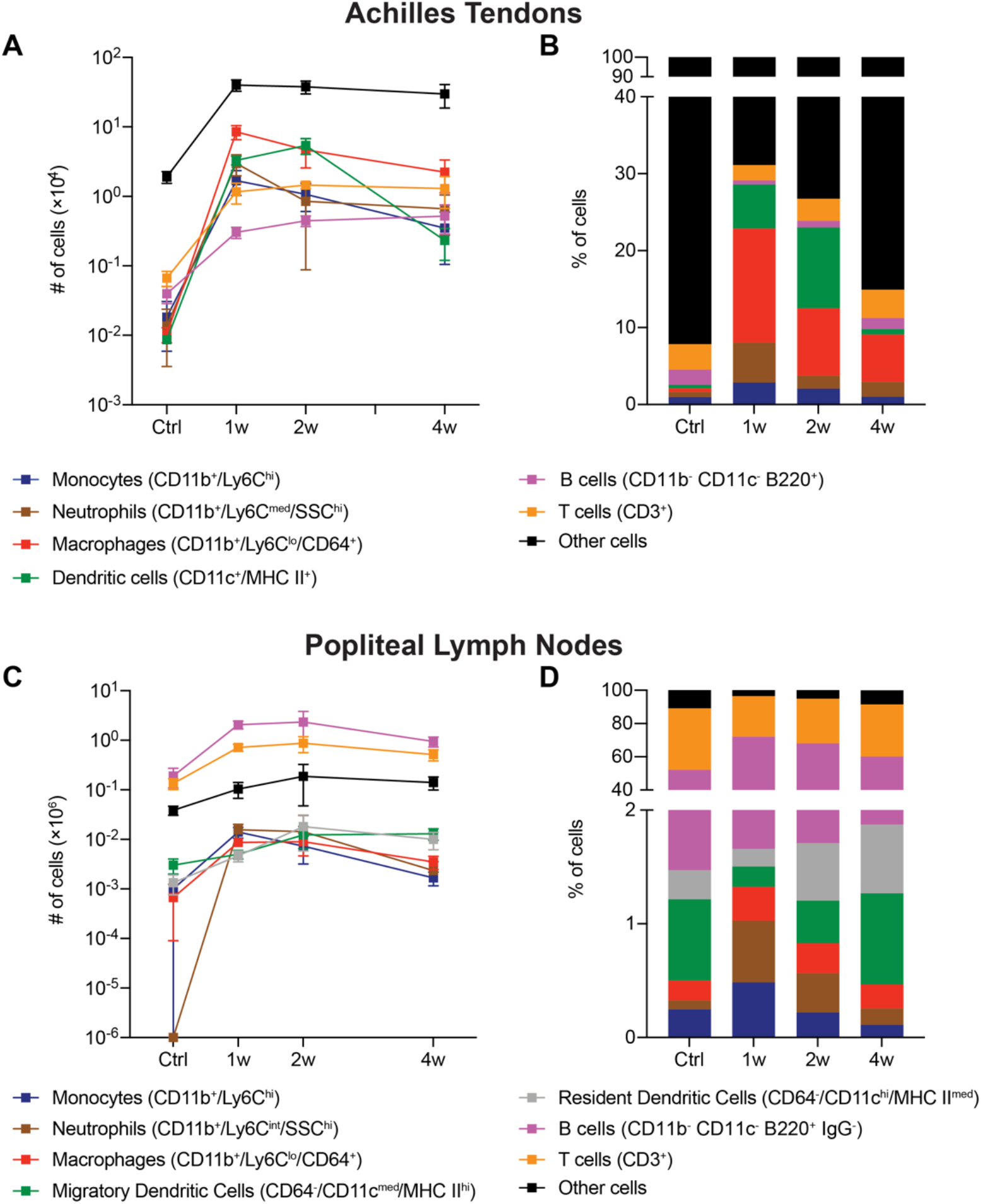
Summary of immune cell changes in Achilles tendons and lymph nodes after Achilles tenotomy and repair. (A) The absolute number of and (B) percentage of monocytes (CD11b^+^/Ly6C^hi^), neutrophils (CD11b^+^/Ly6C^med^/SSC^hi^), macrophages (CD11b^+^/Ly6C^lo^/CD64^+^), dendritic cells (CD11c^+^/MHC II^+^), B cells (CD11b^−^/CD11c^−^/B220^+^), CD4^+^ T cells (CD3^+^/CD4^+^/CD8^−^), CD8^+^ T cells (CD3^+^/CD4^−^/CD8^+^), and other cells from Achilles tendons in control tendons, and tendons either 1 week (1w), 2 weeks (2w), or 4 weeks (4w) after tenotomy and repair. (C) The absolute number of and (D) percentage of monocytes (CD11b^+^/Ly6C^hi^), neutrophils (CD11b^+^/Ly6C^med^/SSC^hi^), macrophages (CD11b^+^/Ly6C^lo^/CD64^+^), migratory dendritic cells (CD11c^int-hi^/MHC II^+^), resident dendritic cells (CD11c^hi^/MHC II^int^), plasma cells (CD11b’/CD11c’/B220’/IgG^+^), B cells (CD11b^−^/CD11c^−^/B220^+^/IgG^−^), CD4^+^ T cells (CD3^+^/CD4^+^/CD8^−^), CD8^+^ T cells (CD3^+^/CD4^−^/CD8^+^), and other cells from popliteal lymph nodes in control tendons, and tendons either 1w, 2w, or 4w after tenotomy and repair.

## Discussion

Changes in innate immune cell populations have been well-studied in tendon injuries(19, 30, 33) but less is known about adaptive immune cell responses. In the current study we sought to characterize the adaptive immune cell response in tendons and tendon-draining lymph nodes to tendon injury and repair. In addition to corroborating innate immune cell data from previous tenotomy studies, we identified a robust adaptive immune cell response after tenotomy in both the tendon and the tendon-draining popliteal lymph node that generally peaks following the innate immune cell response. Maximal T and B cell accumulation in the popliteal lymph node occurred slightly earlier than in the tendon, consistent with the paradigm that T and B cells are initially stimulated and activated by antigen flowing from the affected tissue and then traffic to the affected tissue to carry out their effector functions (6). These results provide novel insight into the facets of the immune system that are involved in the response to tendon injury, and provide a basis for future studies to interrogate their function in tendon injury and healing, with the long term goal of potentially manipulating adaptive immunity to improve tendon healing.

Using a similar Achilles tenotomy and repair model and time points in rats, we previously demonstrated that M1 macrophages accumulate within one week of tendon injury, while M2 macrophages start appearing in appreciable numbers by the second week (30). Although we did not sort macrophages into subtypes, the results in the current study generally follow similar trends in markers of macrophage abundance that we observed in rats (30). Supporting our observations from flow cytometry, the monocyte attractant gene *Ccl2* (12, 17) was also upregulated one week after injury, and the M2 marker *CD206* (30) was elevated at two weeks. Macrophages also appear to directly modulate tendon healing. Depleting macrophages reduces markers of inflammation and improves mechanical properties of tendons at one and two weeks after tendon injury (16), although since M2 macrophages do not begin to accumulate until two weeks after injury, it is possible that an early reduction in inflammation may result in incomplete healing and reduced long-term functional outcomes. Much still remains to be determined about how different macrophage populations interact with other cell types in tendon to modulate tissue repair.

The adaptive immune system is known to play an important role in tissue pathology in disease states, but its role in healing of various musculoskeletal injuries is an emerging area of study(17), and even less is known about lymphocytes in tendon healing. There is also cross-talk between the innate and adaptive immune systems during tissue repair (12). A previous study in rats subjected to an Achilles tendon tear with or without Botox treatment evaluated macrophage and T cell changes through ten days after injury, and identified a response from adaptive immune cells, but conclusions from this study are limited by lack of data on absolute cell numbers, the absence of analysis of uninjured animals, and the lack of statistical analysis of the effect of time on cell accumulation (2).

In the current study, we observed the presence of lymphatic vasculature in the superficial regions of tendon as indicated by the presence of the lymphatic markers LYVE1 and podoplanin (14), similar to previous observations in rat Achilles tendons (32). We also report a small population of tissue resident adaptive immune cells, as well as an induction of *Cxcl12* expression which promotes circulating lymphocyte chemotaxis and tissue entry (12, 17). Based on results from the current study, previous studies evaluating the innate immune response in acutely injured tendons (16, 19, 30, 33), and our understanding of the regulation of adaptive immune cells in other tissue types (3, 12, 17, 18, 25, 29), we have developed a model of immune cell population changes in the context of healing following a tendon tear (Figure 7). Upon injury, there is an accumulation of neutrophils which initiate the inflammatory response and begin to recruit circulating monocytes to the tendon. Monocytes can then differentiate into macrophages and dendritic cells, both of which are able to process antigen from the injured site and present antigen fragments to lymphocytes. The macrophage response through two weeks after injury is dominated by a population of proinflammatory M1 macrophages, which are gradually replaced by an antiinflammatory and tissue regenerative M2 macrophage population. Dendritic cells, which are stimulated by local inflammation, begin to migrate to the draining lymph node within days where they prime naïve CD4^+^ and CD8^+^ T cells. In addition, even prior to high quantity dendritic cell migration from the tendon, soluble antigens and cytokines from the injured tendon flow via the lymphatics to the draining lymph node which contributes to the rapid accumulation of T and B cells. The activated effector CD4^+^ and CD8^+^ T cells then leave the lymph node and migrate to the tendon where they likely help to orchestrate the resolution of inflammation and tissue repair through coordination of tenocyte and other cell activities. Antibody-secreting plasma cells, differentiated from naïve B cells in the lymph node, may also migrate to the tendon to modulate inflammation or repair. Further studies are necessary to more precisely identify the roles of the different lymphocyte populations and the role of the draining lymph nodes in tendon healing.

**Figure 7.**
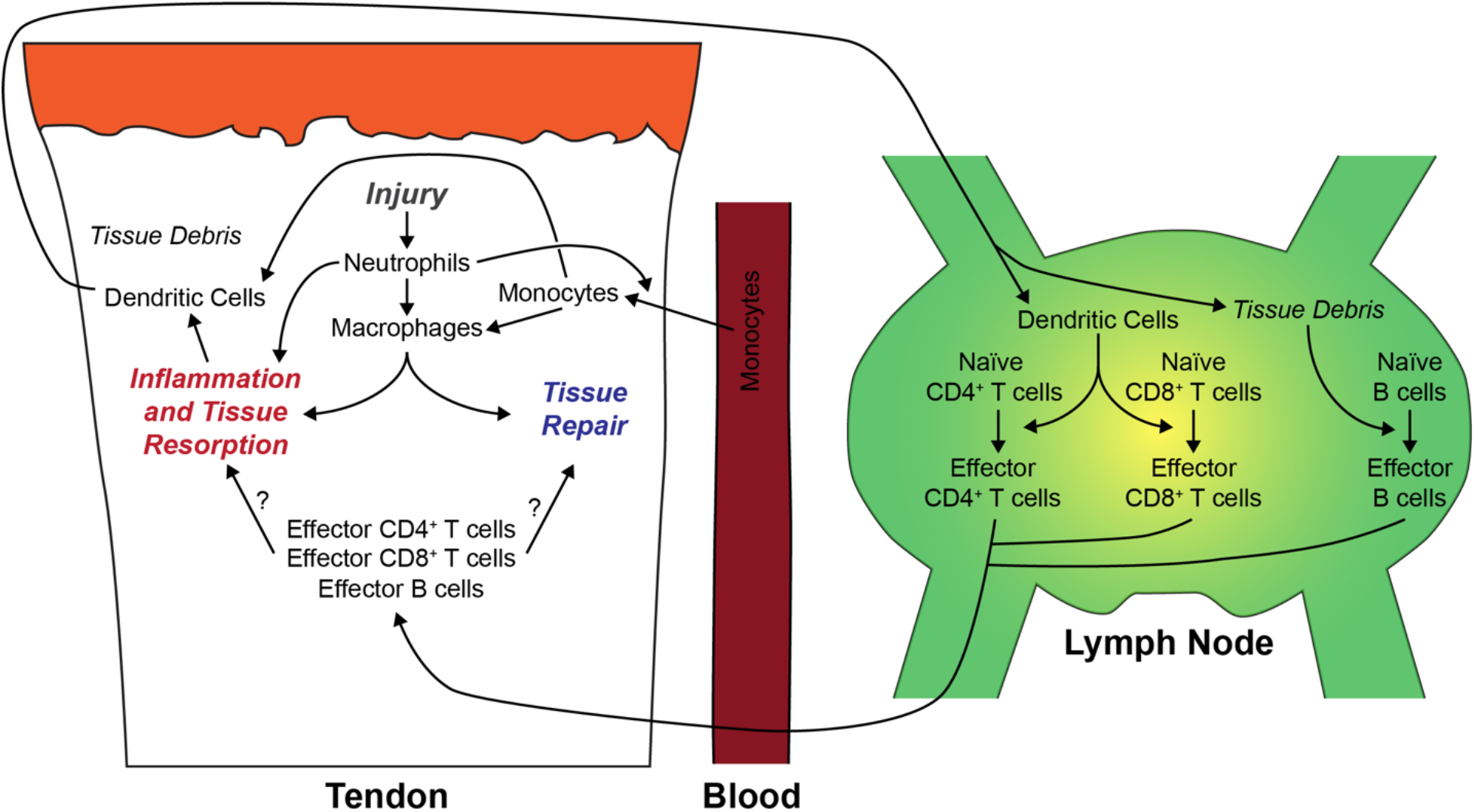
Summary of immune cell changes after tendon injury. An overview of the proposed changes in immune cell abundance and interactions in the context of tendon injury and repair.

There are several limitations to this study. We only evaluated through four weeks after injury, and we expect immune cells continue to play a role in tissue inflammation and repair beyond this point. There are likely also more pronounced changes in early innate immune cells, such as neutrophils, that occur earlier than one week. Although we measured total macrophage abundance after tenotomy and repair, we did not assess M1 and M2 macrophage markers, but did measure the M2 marker CD206 by qPCR. We only evaluated changes in young adult mice, and the immune response is likely to differ as animals age. There are known to be sex-based differences in immune cell biology (26), and as we only included male mice in this study, additional experiments evaluating the role of sex will likely provide additional insight into the biology of tendon repair. Our study focused on the popliteal lymph nodes as these are in closest proximity to the site of injury. However, it is likely that the inguinal and paraaortic lymph nodes, which drain the popliteal lymph node (10), are also involved in the response to a tendon injury. We also did not evaluate changes in tendon mechanics, and integrating this data in the context of changes in immune cell populations could inform the functional role of the innate and adaptive immune systems in tendon healing. Despite these limitations, we feel that this study provides novel insight into the adaptive immune system during tendon healing.

Tendon tears remain a challenging clinical condition (20, 33). Even with improvements in surgical interventions and rehabilitation protocols, many patients continue to have impaired mobility, and up to a third of athletes are unable to return to play after tendon rupture (13, 33, 34). Regenerative strategies for the management of chronic tendon injuries have mostly focused on the delivery of stem cells and growth factors to sites of injured tendon tissue to begin the healing process (4, 8). As such, successful tendon repair requires a fundamental understanding of the inflammatory cascade within musculoskeletal tissue in response to injury. The formation of new tendon tissue relies on a complex interplay between the infiltrating immune cell populations in the epitenon and the resident tenocytes. While the utilization of stem cells and growth factors has been associated with early promising results in animal studies (22), the main limitation of this treatment approach is the bulk administration of these biological agents, without consideration of normal spatial and temporal variations within the acute inflammatory response. In the current study, the presence of adaptive immune cells such as T and B cells was demonstrated at the repair site of an injured Achilles tendon at least one month after the tenotomy and repair was performed. This finding is consistent with the persistent inflammatory cell infiltrates seen on biopsies taken from the Achilles tendons of symptomatic patients with chronic tendinopathy several months after the initial injury (21). Little is known about the contribution of T and B cells to the progression of chronic tendon injuries, but since the immune system plays a critical role in the repair of many different tissue types throughout the body, targeting components of the adaptive immune response and the downstream signaling pathways should be considered in any regenerative therapy for tendon injuries. This highlights the need for additional experiments focused on depletion of adaptive immune cells or inhibition of their inflammatory cytokines to determine the extent in which tendon healing can be augmented.

## Acknowledgements

This work was supported by NIH grants R01-AR063649 and R01-AI079178. The authors have no financial conflicts of interest. The ScxGFP mice in this report were kindly provided by Dr. Ronen Schweitzer of Shriners Childrens Hospital of Portland. We would like to acknowledge technical assistance from Mr. Alex Piacentini at the Hospital for Special Surgery, and useful discussions with Dr. Dragos Dasoveanu, and Dr. Lionel Ivashkiv at the Hospital for Special Surgery.

## Author Contribution Statement

ACN, TML, LMM, SW, KBS, SAR, TTL, and CLM conceived and designed research; ACN, TML, LMM, SW, JBS, NPD performed experiments; ACN, TML, LMM, JBS, NPD, KBS, SAR, TTL, and CLM analyzed data and interpreted results of experiments; ACN, TML, KBS, TTL, and CLM prepared figures, drafted the manuscript, and edited and revised the final manuscript; All authors approved of the submitted and final versions.

